# Enhancing precision medicine for patients with breast cancer brain metastases through functional drug testing

**DOI:** 10.64898/2026.04.18.719400

**Authors:** Ellen Wold, Nathan Merrill, Habib Serhan, Aaron Udager, Chia Jen Liu, Ning Gu, Liwei Bao, Zhaoping Qin, Xu Cheng, Peter Ulintz, Jason Heth, Matthew Soellner, Sofia D. Merajver, Aki Morikawa

## Abstract

Patient-derived organoids from breast cancer brain metastases enable real-time drug sensitivity testing integrated with genomic profiling. Drug response varied by subtype and molecular alterations. PI3K inhibitors showed activity regardless of PIK3CA mutation status. Pronounced tumor heterogeneity highlighted the urgent need for effective therapies personalized for each patient. Functional assays and molecular matching can help tailor therapy for patients who need the most effective next treatment quickly and warrant further translational evaluation to address this unmet need.

## Background

Breast cancer is the second most common malignancy that metastasizes to the brain. Despite surgery and radiation, 30–76% of patients progress within one year. With advances in targeted and antibody-based therapies, systemic treatment has become central to breast cancer brain metastases (BCBrMet) management [1].

Molecular profiling shows that about 50% of BCBrMet harbor potentially actionable genomic alterations.[2]. Molecular-matched therapies may benefit BCBrMet patients, who were previously excluded from clinical trials. Ongoing trials, including Alliance A071701 and ASCO TAPUR, now include these patients [3–4]. However, earlier trials in non-BrMet populations showed limited benefit [3,5], underscoring the need for better molecular cellular guided strategies to select personalized therapies.

Patient-derived xenografts and organoids more accurately capture tumor heterogeneity than cell lines, improving drug sensitivity prediction [6–7]. Particularly, patient-derived organoids (PDOs) preserve tumor microenvironment and tumor cell heterogeneity features that are conducive to clinically relevant drug testing. [8]. In this study, we evaluated the feasibility of real-time, patient-specific drug testing using PDOs developed from resected BCBrMet, integrating functional assays with genomic profiling.

## Results

### Cohort Characteristics

Between September 2019 and November 2024, 29 BCBrMet surgical samples were collected from 27 patients. PDOs were established in 21 samples from 19 patients. PDOs were not generated due to insufficient viable cells, primary brain tumor, or logistic constraints (including pandemic-related disruptions). Two patients provided multiple samples: BC25 had serial frontal lobe resections, and BC26 had synchronous cerebellar and occipital resections. Median age at BCBrMet resection was 55 years (range 30–72), with median ages at initial, metastatic, and BCBrMet diagnosis of 47, 54, and 55 years, respectively.

Primary tumor subtypes included HR+/HER2+ (37%), HR−/HER2+ (21%), HR+/HER2− (16%), and HR−/HER2− (26%). Subtype distribution for BrMet was HR+/HER2+ (21%), HR−/HER2+ (32%), HR+/HER2− (21%), and HR−/HER2− (26%). Most patients received pre-resection systemic therapy (neoadjuvant/adjuvant 68%, metastatic 68%). Prior CNS therapies included surgery (11%), stereotactic radiosurgery (SRS) (26%), and whole-brain radiation therapy (WBRT) (5%). Postoperative survival at 1, 6, and 12 months was 16%, 21%, and 37%, respectively (Table 1, Supplement 1).

**Table 1.** Clinical and pathological characteristics of cohort. Data is presented as number (%) unless otherwise noted. Abbreviations: ER, estrogen receptor; PR, progesterone receptor; HER2, human epidermal growth factor receptor 2; HR, hormone receptor; CNS, central nervous system.

| All patients (n=19) |  |
| --- | --- |
| **Age (years)** |  |
| *Median, (range)* |  |
| At breast cancer diagnosis | 47 (28-66) |
| At metastatic disease diagnosis | 54 (30-66) |
| At brain metastasis diagnosis | 55 (30-72) |
| At brain metastasis resection | 55 (30-72) |
| **Brain Metastasis Subtype** |  |
| HR+/HER2+ | 4 (21%) |
| HR-/HER2+ | 6 (32%) |
| HR+/HER2- | 4 (21%) |
| HR-/HER2 - | 5 (26%) |
| **Primary Tumor Subtype** |  |
| HR+/HER2+ | 7 (37%) |
| HR-/HER2+ | 4 (21%) |
| HR+/HER2- | 3 (16%) |
| HR-/HER2 - | 5 (26%) |
| **Sites of Metastasis** |  |
| Liver | 7 (37%) |
| Lung | 11 (58%) |
| Lymph Node | 15 (79%) |
| Bone | 11 (58%) |
| Leptomeninges | 3 (16%) |
| **Prior Systemic Therapy** |  |
| Neoadjuvant/adjuvant | 13 (68%) |
| Metastatic systemic therapy | 13 (68%) |
| Immunotherapy | 4 (21%) |
| **Prior CNS Treatment** |  |
| Surgical Resections | 2 (11%) |
| Stereotactic Radiosurgery | 5 (26%) |
| Whole Brain Radiation Therapy | 1 (5%) |
| CNS directed systemic therapy | 6 (32%) |
| **Survival After Resection** |  |
| Less than 1 month | 3 (16%) |
| Less than 6 months | 4 (21%) |
| Less than 12 months | 7 (37%) |

### Molecular Characterization

RNA profiling was performed on 14 cases with sufficient material for molecular characterization, at sample collection to guide drug selection. Expression data demonstrated high AKT1 expression some samples, which informed drug choices. The full listing of genes from RNA profiling is in Supplement 2.

Retrospective targeted mutational analysis was conducted on all PDOs to assess correlations with experimental drug response data. TP53 mutations were present in 81%, PIK3CA in 24%, and other mutations (ESR1, ERBB2, BRCA1, PMS1, RAD51) ranged from 5–10%. Gene expression profiling for ERBB2 showed large variability of expression (ERBB2: 5.63–15.46). ERBB2 levels generally aligned with HER2 status, but some HER2−negative cases (e.g., BC32) exhibited relatively high ERBB2 expression (Supplement 3). The initial and recurrent lesions (BC25-1 and BC25-2) shared similar variants (Figure 1), whereas the synchronous lesions (BC26-1 and BC26-2) differed in PIK3CA mutation status (Figure 2).

**Figure 1.**
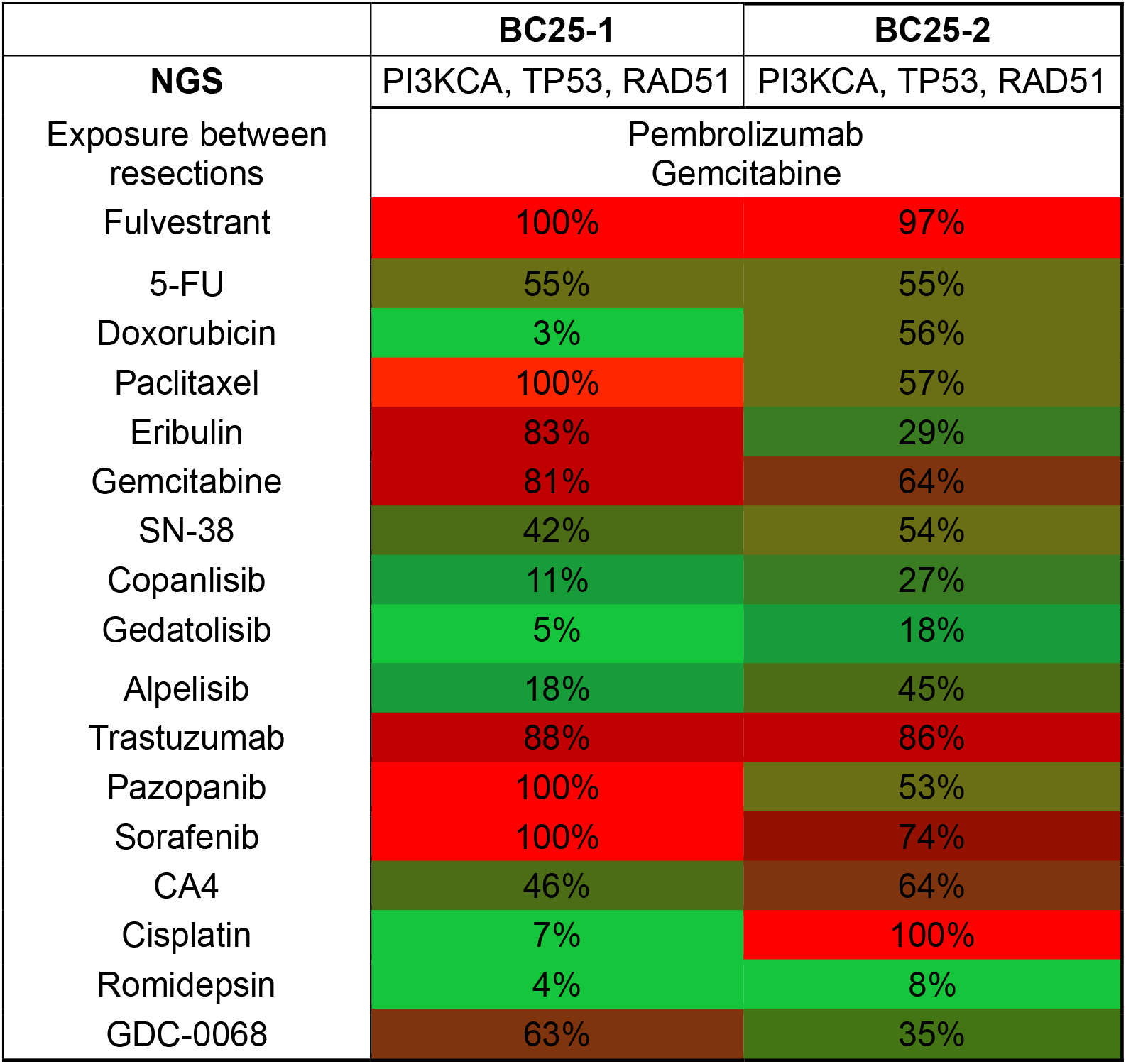
Molecular characteristics and drug sensitivity profiles of sequential right frontal lobe metastasis resections. This figure presents the molecular alterations, interim clinical exposures, and drug sensitivity results for two sequential right frontal lobe metastasis resections in a patient with TNBC. The table demonstrates percent viability assessed by drug assay, where lower values indicate greater sensitivity. Abbreviations: BC25-1 and BC25-2 represent the first and second sequential resections, respectively. NGS, next-generation sequencing; TNBC, triple-negative breast cancer; BC25-1 and BC25-2 represent the first and second sequential resections, respectively.

**Figure 2.**
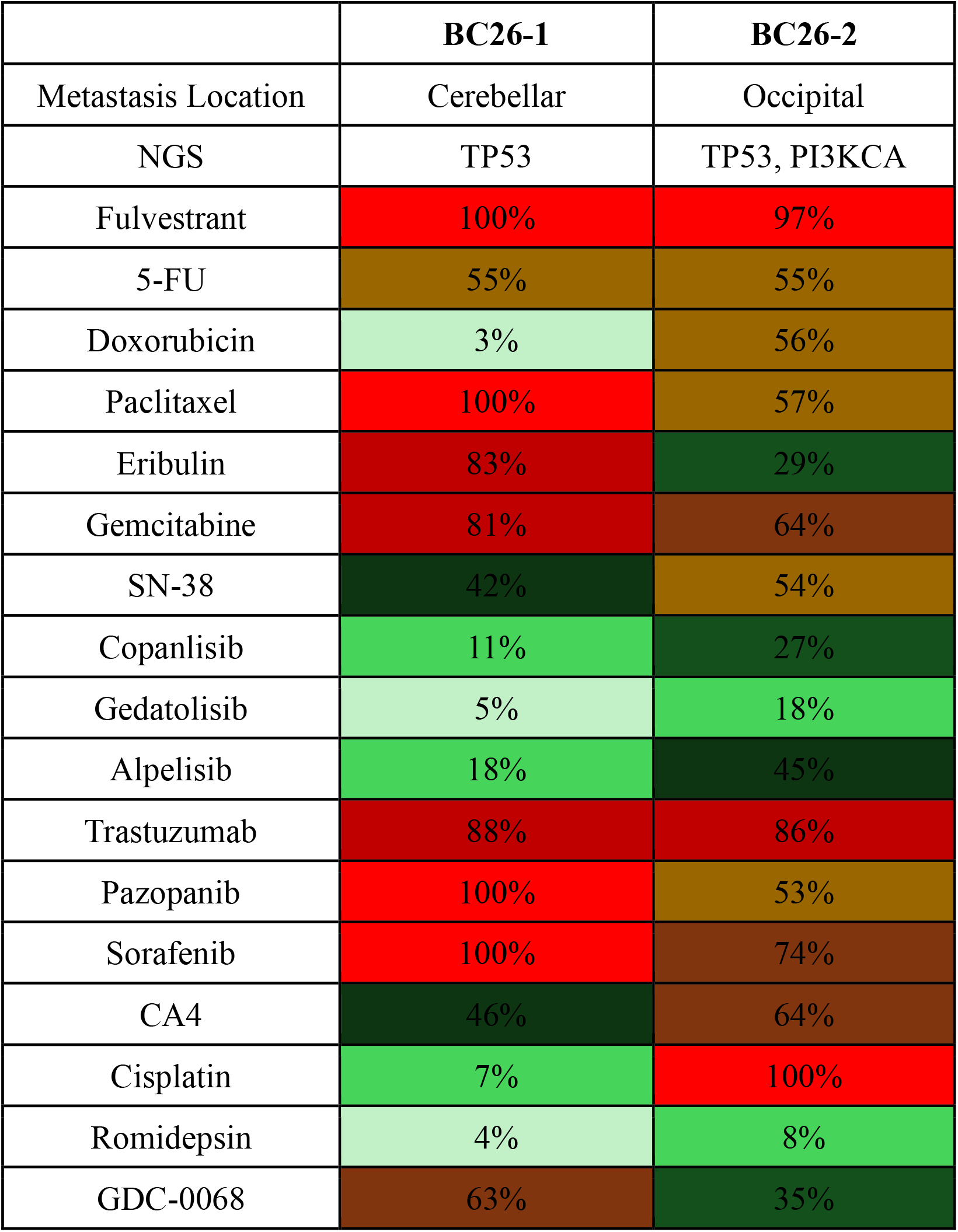
Drug sensitivity profiles and genomic features of simultaneous cerebellar and occipital metastases. Figure presents metastatic locations, genomic alterations, and drug sensitivity results for simultaneous cerebellar (BC26-1) and occipital (BC26-2) brain metastases resected from a patient with triple-negative breast cancer. Percent viability assessed by drug assay is shown; lower values indicate greater sensitivity. Abbreviations: BC26-1 and BC26-2 represent cerebellar and occipital metastases, respectively. NGS, next-generation sequencing; TNBC, triple-negative breast cancer

Significant CNVs are summarized in Supplement 3. Briefly, recurrent events involved alterations on Chr1, Chr8, Chr11, and Chr17, with additional clusters of ChrX deletions in TNBC cases. PIK3CA amplification was observed in BC18, BC26-2, BC27, and BC38. BC26-2 also harbored a PIK3CA mutation not present in the synchronous lesion (BC26-1).

### Drug Testing Data

Drug-sensitivity testing was performed on 20 of 21 PDOs; one lacked sufficient viable tissue. Median interval from resection to results was 13 days (range: 7–22 days). 55 agents, including chemotherapies and targeted drugs, were tested (Supplement 4). Each PDO was exposed to a median of 13 agents (range: 1–21). Drug selection was guided by prior treatments and clinical options for further therapies. Next-generation sequencing (NGS) data were available for 16 patients (84%) at resection. Selected illustrative cases are discussed below.

#### PI3K Pathway Inhibitors *(Copanlisib, Gedatolisib, Alpelisib, Paxalisib, GDC-0068)* show sensitivity even after multiple lines of therapy, but levels differ between agents

The PI3K pathway plays a key role in BCBrMet development [2]. Pre-drug testing RNA expression results showed elevated *AKT1* expression (Supplement 2), leading to the use of PI3K inhibitors regardless of *PIK3CA* mutation status. PI3K inhibitors produced heterogeneous responses across PDOs, with sensitivity levels being independent of PIK3CA mutation status (Supplement 4). Drug screening used Cmax (maximum plasma concentration) from FDA labels and clinical trials.

Of five PDOs with *PIK3CA* mutations, four demonstrated moderate to high sensitivity to PI3K inhibitors (Figure 3). Outcomes were poor – all four patients died within one year of resection, including two within one month. None received PI3K inhibitors regardless of subtype (Table 2). For example, BC29 (HR+/HER2+, PI3K mutation+) received multiple therapies tested in PDO assays, including paclitaxel (C-max 76.9), trastuzumab (C-max 96.4), and tucatinib (C-max 90.9). She died 10 months after operation. PDO screening, completed in 16 days postoperatively, suggested potential for benefit from PI3K inhibitors, notably paxalisib (C-max 14) (Figure 3, Table 2).

**Table 2.**
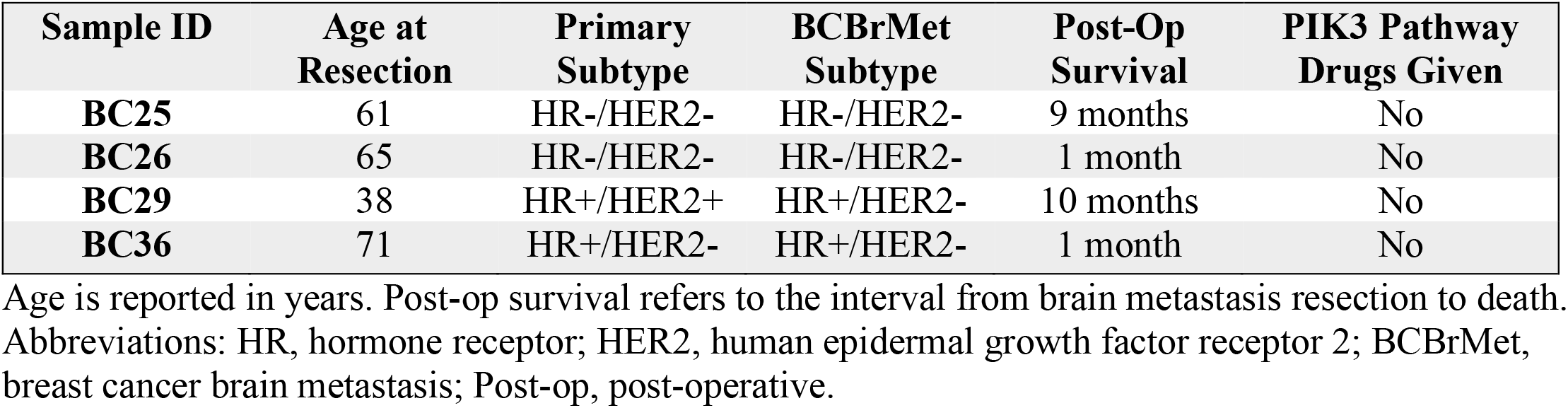
Clinical characteristics and outcomes of patients with PI3K pathway mutations.

| **Sample ID** | **Age at Resection** | **Primary Subtype** | **BCBrMet Subtype** | **Post-Op Survival** | **PIK3 Pathway Drugs Given** |
| --- | --- | --- | --- | --- | --- |
| **BC25** | 61 | HR-/HER2- | HR-/HER2- | 9 months | No |
| **BC26** | 65 | HR-/HER2- | HR-/HER2- | 1 month | No |
| **BC29** | 38 | HR+/HER2+ | HR+/HER2- | 10 months | No |
| **BC36** | 71 | HR+/HER2- | HR+/HER2- | 1 month | No |

Age is reported in years. Post-op survival refers to the interval from brain metastasis resection to death. Abbreviations: HR, hormone receptor; HER2, human epidermal growth factor receptor 2; BCBrMet, breast cancer brain metastasis; Post-op, post-operative.

**Figure 3.**
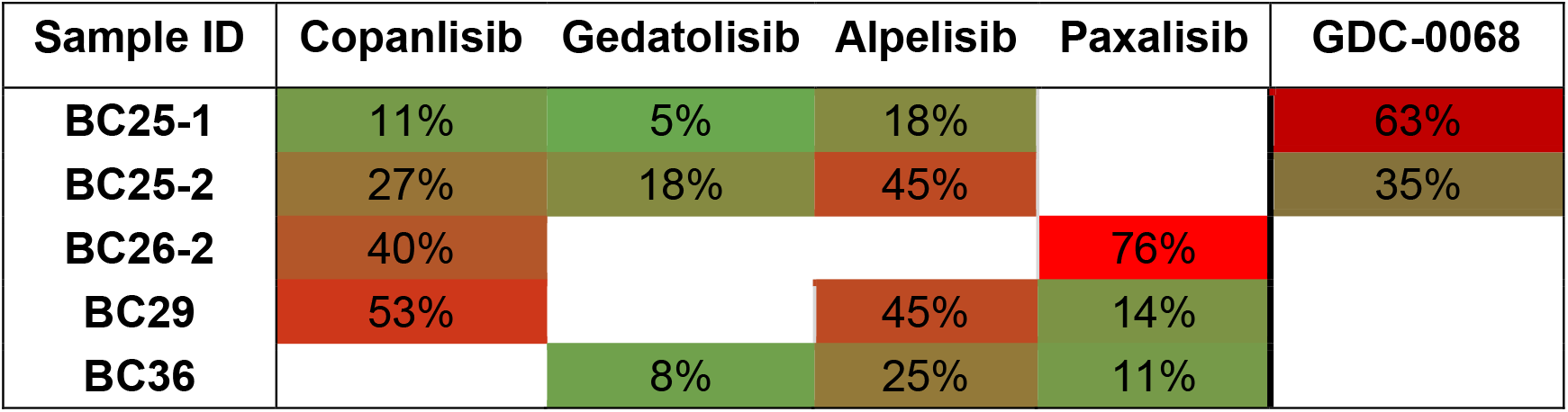
Comparative cell viability of PIK3CA-mutated PDOs exposed to PI3K inhibitors at C-max. PDOs with PI3KCA mutations were treated with copanlisib, gedatolisib, alpelisib, paxalisib, and GDC-0068 at concentrations corresponding to C-max for each drug. Values represent the percentage of viable cells relative to untreated controls as determined by CellTiter-Glo 3D, with lower values indicating higher sensitivity. Blank cells indicate the treatment was not performed for the corresponding PDO. Abbreviations: PDO, patient-derived organoid; C-max, maximum clinical plasma concentration.

#### BC25: Drug sensitivity changes post-immunotherapy and radiation exposure

BC-25 was a 61-year-old Caucasian female with TNBC who underwent sequential resections of a right frontal lobe metastasis. Before initial resection, she received adjuvant chemotherapy (doxorubicin, cyclophosphamide, nab-paclitaxel, capecitabine, and bevacizumab) followed by gemcitabine, cisplatin, paclitaxel, and eribulin for metastatic disease. In the six months between resections, treatment included pembrolizumab, gemcitabine, and brain radiation (Supplement 1). DNA profiling demonstrated similarity between the 2 specimens (Figure 1), but CNV differed in *PRCACA, SDHD, MYC, RECQL4*, and *FGFR1* (Supplement 3). Drug sensitivity changed after exposure to immunotherapy and brain radiation Notably, BC25-2 showed reduced response to PI3K inhibitors, except GDC-0068 (Figure 1).

#### BC26: Synchronous lesions demonstrate tumor heterogeneity with differences in drug sensitivity

BC-26 was a 65-year-old African American female with TNBC who underwent simultaneous resection of cerebellar (BC26-1) and occipital (BC26-2) metastases following nab-paclitaxel and WBRT (Supplement 1). Lesions differed in PIK3CA mutation status (BC26-1 negative, BC26-2 positive). However, BC26-1 was more responsive to all drugs tested, including PI3K inhibitors (Figure 2). RNA analysis revealed that BC26-2 had increased PIK3CA gene expression and CNV, which were not present in BC26-1. (Supplement 2, 3).

## Discussion

PDO drug testing enhances precision medicine through complementary functional assays [8]. We collected BCBrMet from clinically indicated resections to evaluate real-time drug testing feasibility and assess multiomic molecular complexity is correlated to drug response. PDOs were established and tested in most cases despite pandemic-related challenges, except for a few cases with essentially no viable cells. Drug testing was completed within a median of 13 days, a clinically meaningful timeframe for potentially contributing to treatment decisions in advanced cancer.

PDOs were mainly HER2−positive and triple-negative (Table 1, Supplement 1), reflecting the higher incidence of BrMet in these subtypes [1], though we also analyzed rarer HR+/HER2+ cases. Mutation profiles frequently showed TP53 and PI3K/AKT pathway alterations as previously reported [9] (Supplement 3). Targeted RNA panels revealed CNV in genes associated with immune/inflammatory pathways and upregulation of extracellular matrix, similar to the finding from our previous published BCBrMet study (Supplement 2) [7]. Modulation of these pathways can be further examined to overcome potential drug resistance in BCBrMet.

BCBrMet has been associated with the PI3K/AKT pathway [9]. We found significant activity, largely independent of PIK3CA mutation status, highlighting the potential added value of functional testing (Supplement 4). Notably, in synchronous lesions, the PIK3CA-mutant sample was less responsive to PI3K inhibitors and had CN amplification, suggesting reduced sensitivity may be due to co-occurring alterations or non-mutational signaling alterations (Figure 2, Supplement 2). Further mechanistic investigations are warranted to determine the impact of the CNV on PI3K pathway inhibition. In pre- and post-immunotherapy/radiation PDO pairs, we observed differences in CNV rather than new DNA mutations (Supplement 2, 3). Others have reported associations between the genes (e.g. CD274, PRKACA, SDHD, RECQL4) with CNV and immunotherapy response, suggesting the potential of specific CNV as biomarkers [10–13].

These findings underscore the molecular heterogeneity of BCBrMet and the difficulty of predicting drug response from DNA/RNA analyses when multiple levels of molecular alterations exist. This complexity contributes to the varied and mixed treatment outcomes seen in metastatic cancer. In current clinical practice, the molecular matching relies heavily on the detection of DNA mutations to select drug therapy, in spite of poor reliability of such guidance to select drugs in clinical trials [3,5]. Our study suggests complementary approaches, such as RNA analysis and functional drug testing, may help guide therapy and uncover non mutational compensatory mechanisms that drive drug resistance. This is especially important in BCBrMet, because the opportunity for multiple drug regimens after progression beyond standard of care is highly limited. Thus, understanding what drugs the brain metastatic lesions are resistant to stands to spare patients from exposure to potentially ineffective therapies.

This study is limited by a small sample size and restricted access to newer agents like antibody-drug conjugates (ADCs) and a lack of active and proliferative immune components in PDOs. However, given the uniqueness of PDOs from BCBrMet, our findings provide valuable insights. We have since acquired ADCs and are developing enhanced PDO platforms for immunotherapy. The platform is now optimized for future clinical use. In our cohorts, correlations between the drug testing and prospective patient outcome with the tested drugs were scarce due to limited survival and drug availability (in our drug panel, as mentioned, or no clinical labeling indication), and because we often lacked advanced knowledge of the next prescribed drug for panel inclusion. Based on the work described here, the new platform will be aimed at performing functional drug testing in parallel, without informing treatment, for patients on clinical trials, to establish the platform’s accuracy in predicting sensitivity and resistance.

Given the rarity and small size of these samples, we limited long-term PDO expansion to preserve clinical characteristics and avoid altering drug responses. When enough cells were available, we established cell lines and PDX models for further investigations [7,14].

## Conclusion

Real-time, patient-tailored drug testing in 3-dimensional, minimally cultured organoids that preserve the tumor microenvironment, as our group has previously demonstrated in bladder cancer [15], is feasible for BCBrMet patients undergoing resection. Molecular profiling alone appears inadequate for guiding targeted therapy; however, complementary functional testing may help optimize treatment choice.

## Methods

### Patient Enrollment and PDO Establishment

Patients undergoing clinically indicated biopsy or resection of brain metastases from solid tumors provided written informed consent under a University of Michigan IRB–approved CNS repository tumor bank protocol (HUM00024610). PDO and PDX models were generated under an IRB-approved protocol (HUM00103662). The study was conducted in accordance with the Declaration of Helsinki and applicable regulations, and specimens were deidentified prior to laboratory receipt. Eligibility criteria for the PDO study included: informed consent to the CNS tissue collection protocol, diagnosis of invasive breast cancer, confirmed brain lesion requiring surgery or biopsy, availability of tumor tissue remaining after clinical pathology review, and pathological confirmation that malignant cells were consistent with breast origin.

Fresh tumor tissue was collected intraoperatively and immediately transferred on ice to the laboratory. Tumors were minced into small fragments and enzymatically dissociated using the MACS Tumor Dissociation Kit (Miltenyi Biotec), with mechanical agitation and incubation at 37°C for up to 60 minutes. The resulting suspension was filtered through a 70-μm cell strainer to generate a single-cell suspension. PDO culture was performed using a three-dimensional culture system composed of DMEM (Thermo Fisher Scientific), supplemented with 10% fetal bovine serum (Thermo Fisher Scientific), 1% Matrigel (Corning), B-27 supplement (Thermo Fisher Scientific), antibiotic-antimycotic solution (Thermo Fisher Scientific), and gentamicin (Thermo Fisher Scientific). The media was additionally enriched with recombinant human growth factors including EGF (25 µg/500 ml; Sigma Aldrich), Heregulin β-1 (25 µg/500 ml; Stemcell Technologies), KGF/FGF-7 (5 µg/500 ml; Stemcell Technologies), FGF-10 (5 µg/500 ml; Stemcell Technologies), Noggin (50 µg/500 ml; Stemcell Technologies), and RSPO1 (250 µg/500 ml; Stemcell Technologies). Cells were plated onto ultra-low attachment culture dishes [15]. Organoids were maintained at 37°C in a humidified incubator with 10% CO_2_, with medium changes every 2–3 days.

### Molecular Profiling and Next-Generation Sequencing

Targeted DNA sequencing was conducted on BCBrMet tissue. DNA was extracted, and NGS libraries were constructed and sequenced using an Ion Torrent NGS system. Variants were detected from aligned reads, annotated, filtered, and manually curated to identify prioritized alterations. Gene-based copy number ratios were calculated. Illumina-based RNAseq libraries were constructed, sequenced, and analyzed to quantify gene expression [16–18]. Counts per million (CPM) values were normalized using edgeR and expression was displayed as log2 CPM values. Additionally, RNA samples underwent assessment with the Nanostring BC360 panel, which employs digital molecular barcoding technology to quantify RNA expression levels. The resulting data were processed and interpreted using nSolver software.

### Drug Testing

Established PDOs were plated in 96-well ultra-low attachment plates (3,000-5,000 cells/well, 100 μL/well) and centrifuged (120 × g, 5 min). Organoids were exposed in triplicate to a panel of up to 21 molecularly targeted and standard-of-care drugs in a dose-response format when material was sufficient or at clinically relevant Cmax concentrations when material was limited. After 120 hours, cell viability was measured using CellTiter-Glo 3D (Promega), and drug sensitivity scores (DSS3) were calculated. Drugs were categorized by activity score as “very active,” “active,” or “inactive,” per established thresholds [16, 19]. Each experiment included vehicle negative controls, and all drug testing was performed in at least two technical replicates to validate reproducibility.

A list of drugs was created based on the clinical history, clinically available genomic testing results (usually done on non-brain metastases samples), and Nanostring testing results of the brain metastases at the time of sample collection. A set of commonly used drugs for breast cancer (doxorubicin, paclitaxel, and fulvestrant) was included in most of the cases, regardless of clinical history, when the sample was considered to have enough cells to test multiple drugs. The number of drugs tested per sample were determined based on quantity of cells to achieve a minimum of 3,000 cells per well.

### Correlative Clinical Analysis

A retrospective chart review was performed for PDO source patients to collect data on demographics, primary and brain metastases receptor status, clinically-available genomic testing results, treatment history, and clinical outcomes. These clinical parameters were matched to PDO drug sensitivity data and molecular profiles to evaluate associations between PDO-predicted sensitivities, therapies received, and clinical responses. The University of Michigan’s AI tool U-Maizy (February 2025) was employed to assist with data summarization.

## Author Contributions

Conceptualization: N.M., S.D.M.,and A.M.

Formal analysis: E.W. A.U., N.M., and A.M.

Supervision: N.M., S.D.M., and A.M.

Funding acquisition: S.D.M. and A.M.

Methodology: N.M., J.H., M.S., P.U., S.D.M.,and A.M.

Project administrations: N.M., A.U., J.H., S.D.M., and A.M.

Investigation: E.W., N.M., H.S., A.U., C.J.L., N.G., L.B., Z.Q., X.C., and A.M.

Writing-original draft: E.W., N.M., and A.M.

Writing-review and editing: All authors

## Ethics Declaration

Patients undergoing clinically indicated biopsy or resection of brain metastases from solid tumors provided written informed consent under a University of Michigan IRBMED–approved CNS repository tumor bank protocol (HUM00024610). PDO and PDX models were generated under an IRB-approved protocol (HUM00103662). The study was conducted in accordance with the Declaration of Helsinki and applicable regulations, and specimens were deidentified prior to laboratory receipt.

